# Fast protein structure searching using structure graph embeddings

**DOI:** 10.1101/2022.11.28.518224

**Authors:** Joe G Greener, Kiarash Jamali

## Abstract

Comparing and searching protein structures independent of primary sequence has proved useful for remote homology detection, function annotation and protein classification. Fast and accurate methods to search with structures will be essential to make use of the vast databases that have recently become available, in the same way that fast protein sequence searching underpins much of bioinformatics. We train a simple graph neural network using supervised contrastive learning to learn a low-dimensional embedding of protein structure. The method, called Progres, is available as software at https://github.com/greener-group/progres and as a web server at https://progres.mrc-lmb.cam.ac.uk. It has accuracy comparable to the best current methods and can search the AlphaFold database TED domains in a tenth of a second per query on CPU.

## Introduction

A variety of methods have been developed to compare, align and search with protein structures [1]. Since structure is more conserved than sequence [2] these methods have proved useful in remote homology detection [3], protein classification [4], inferring function from structure [5], clustering large databases [6, 7] and assessing the accuracy of structure predictions. Global coordinate comparisons like TM-align [8] provide interpretable scores that are comparable across protein size, with a challenge being how to align the residues independent of the primary sequence. Mathematical representations of 3D space such as 3D Zernike descriptors [9, 10] avoid this issue but are limited in accuracy. Other approaches include comparing residue-residue distances [11–14], which can access precise geometries conserved in the structural core, and considering local geometry [15]. The highest accuracy methods tend to be careful comparisons based on coordinates like Dali [13], but searching large structural databases such as the AlphaFold Protein Structure Database [16, 17] or the ESM Metagenomic Atlas [18] with these methods is slow. Recently Foldseek [19] has addressed this problem by converting protein structure into a sequence of learned local tertiary motifs. It then uses the rich history of fast sequence searching in bioinformatics to dramatically reduce the pairwise comparison time of the query with each member of the database. It follows that to further reduce search time, the pairwise comparison step should be made even faster.

Inspired by the impressive performance of simple graph neural networks (GNNs) using coordinate information for a variety of molecular tasks [20], we decided to train a model to embed protein structures into a low-dimensional representation. Two embeddings can be compared very quickly by cosine similarity and a query can be compared to each member of a pre-embedded database in a vectorised manner on CPU or GPU. It makes sense to use expertly-curated classifications of protein structures when training such an embedding [21, 4, 22]; we use supervised contrastive learning [23] to allow the embedding to be learned in a manner that reflects such an understanding of protein structure space and returns search results consistent with it.

A number of recent methods have used protein structure graph embeddings [24–26] and contrastive learning [26–28]. Embedding protein folds has also been done using residue-level features [29, 7], and GNNs acting on protein structure have been used for function prediction [30]. Other studies have used unsupervised contrastive learning on protein structures and show that the representations are useful for downstream prediction tasks including protein structural similarity [31–33]. Contrastive learning using protein classifications has also improved language models for protein sequences, showing clustering that better preserves protein structure space [34]. Protein structure has been incorporated into language models more broadly, often with the intention of searching for remote homology [35–40]. Progres provides a fast and accurate alternative to these methods that is available as a web server and as software.

## Results

We trained a simple GNN, called Progres (PROtein GRaph Embedding Search), to embed a protein structure independent of its sequence (see Figure 1A). Since we use distance and torsion angle features based on coordinates the embedding is SE(3)-invariant, i.e. it doesn’t change with translation or rotation of the input structure. As shown in Figure 1B, supervised contrastive learning [23] on SCOPe domains [21, 41] is used to train the model, moving domains closer or further apart in the embedding space depending on whether they are in the same SCOPe family or not. Sinusoidal position encoding [42] is also used to allow the model to effectively use information on the sequence separation of residues. The main intended use of such an embedding is fast searching for similar structures by comparing the embedding of a query structure to the pre-computed embeddings of a database of structures. Our model does not give structural alignments, but if these are required they can be computed with tools like Dali after fast initial filtering with Progres. The impact of changing model hyperparameters is shown in Table S1. The distribution of values across the embedding dimensions is shown in Figure S1.

**Figure 1.**
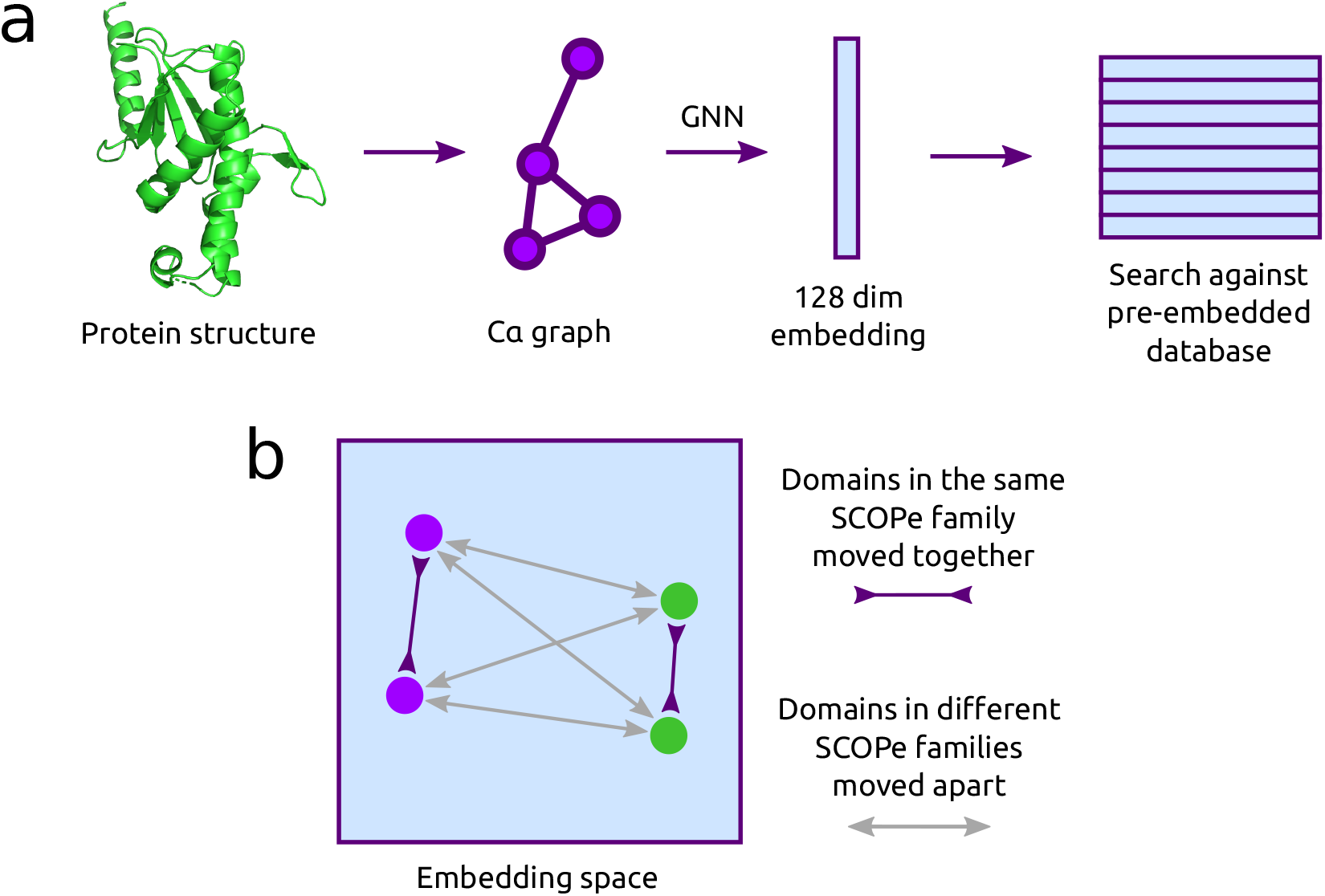
Protein structure embedding. (A) Protein domains are treated as a graph with Cα atoms as nodes and edges between Cα atoms within 10 Å. A GNN embeds the graph into a 128-dimensional representation. This can be compared quickly to a pre-embedded search database. (B) Supervised contrastive learning [23] is used to train the model, with embeddings for domains in the same SCOPe family pushed together and embeddings for domains in different SCOPe families pushed apart. Structures can be automatically split into domains before searching.

In order to assess the accuracy of the model for structure searching, we follow a similar procedure to Foldseek [19]. Since our model is trained on SCOPe domains it is important not to use domains for training that appear in the test set. We select a random set of 400 domains from the Astral 2.08 40% sequence identity set for testing. No domains in the training set have a sequence identity of 30% or more to these 400 domains. This represents the realistic use case that the query structure has not been seen during training - for example it is a predicted or new experimental structure - but other domains in the family may have been seen during training. The easier case of searching with the exact domains used for training gives superior results that are not reported here, and the harder case of searching with completely unseen folds is discussed later.

As shown in Table 1 our model has sensitivity comparable to Dali [13] and Foldseek-TM [19] for recovering domains in SCOPe from the same fold, superfamily and family. Its strong performance at the fold level indicates an ability to find remote homologs. Progres is more sensitive than the EAT [34] and ESM-2 [18] protein language model embeddings, and also the baseline sequence searching method of MMseqs2 [43]. This indicates the benefits of comparing structures rather than just sequences for detecting homology. Figure 2A-C shows the performance across different SCOPe classes, protein sizes and contact orders. Progres does particularly well on all-β domains, smaller domains and domains with higher contact order. This ability to do well in cases where residues separate in sequence form contacts is possibly due to the lack of primary sequence information in the embedding, compared to a method like Foldseek that retains the sequence order for searching. It has lower performance on membrane proteins and larger domains. As shown in Figure 2D, performance drops when the number of embedding dimensions is below 32.

**Table 1.**
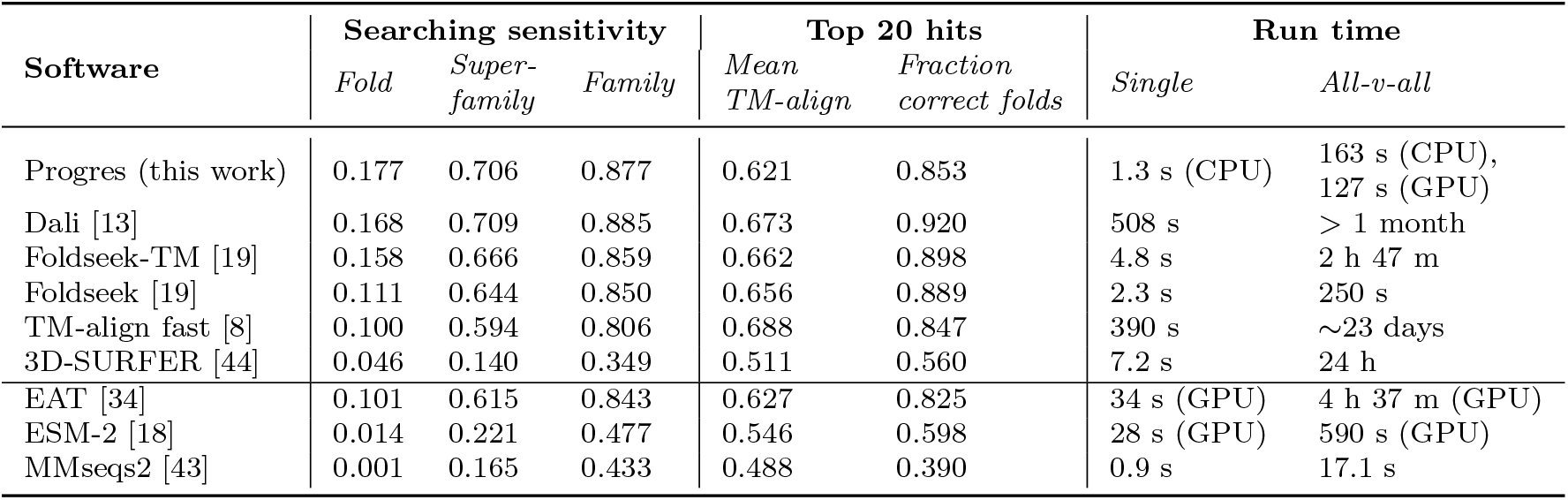
Comparison of ability to retrieve homologous proteins from SCOPe. A similar procedure to Foldseek [19] is followed for searching sensitivity with a set of 400 domains. For each domain the fraction of true positives (TPs) detected up to the first incorrect fold is calculated (higher is better). TPs are same family in the case of family-level recognition, same superfamily and not same family in the case of superfamily-level recognition, and same fold and not same superfamily in the case of fold-level recognition. The mean of this fraction over all 400 domains is reported. Run time (single) is the time taken to search a structure of 150 residues (d1a6ja in PDB format) against all the 15,177 Astral 2.08 40% sequence identity set domains, with the database pre-prepared. Run time (all-v-all) is the time taken to calculate all pairwise distances between the 15,177 domains from structure. EAT, ESM-2 and MMseqs2 use sequence not structure for searching. 3D-SURFER and EAT are trained with structural information and may have seen proteins in the test set during training.

| Software | Searching sensitivity |  |  | Top 20 hits |  | Run time |  |
| --- | --- | --- | --- | --- | --- | --- | --- |
|  | *Fold* | *Super-family* | *Family* | *Mean TM-align* | *Fraction correct folds* | *Single* | *All-v-all* |
| Progres (this work) | 0.177 | 0.706 | 0.877 | 0.621 | 0.853 | 1.3 s (CPU) | 163 s (CPU),<br>127 s (GPU) |
| Dali [13] | 0.168 | 0.709 | 0.885 | 0.673 | 0.920 | 508 s | > 1 month |
| Foldseek-TM [19] | 0.158 | 0.666 | 0.859 | 0.662 | 0.898 | 4.8 s | 2 h 47 m |
| Foldseek [19] | 0.111 | 0.644 | 0.850 | 0.656 | 0.889 | 2.3 s | 250 s |
| TM-align fast [8] | 0.100 | 0.594 | 0.806 | 0.688 | 0.847 | 390 s | ~23 days |
| 3D-SURFER [44] | 0.046 | 0.140 | 0.349 | 0.511 | 0.560 | 7.2 s | 24 h |
| EAT [34] | 0.101 | 0.615 | 0.843 | 0.627 | 0.825 | 34 s (GPU) | 4 h 37 m (GPU) |
| ESM-2 [18] | 0.014 | 0.221 | 0.477 | 0.546 | 0.598 | 28 s (GPU) | 590 s (GPU) |
| MMseqs2 [43] | 0.001 | 0.165 | 0.433 | 0.488 | 0.390 | 0.9 s | 17.1 s |

**Figure 2.**
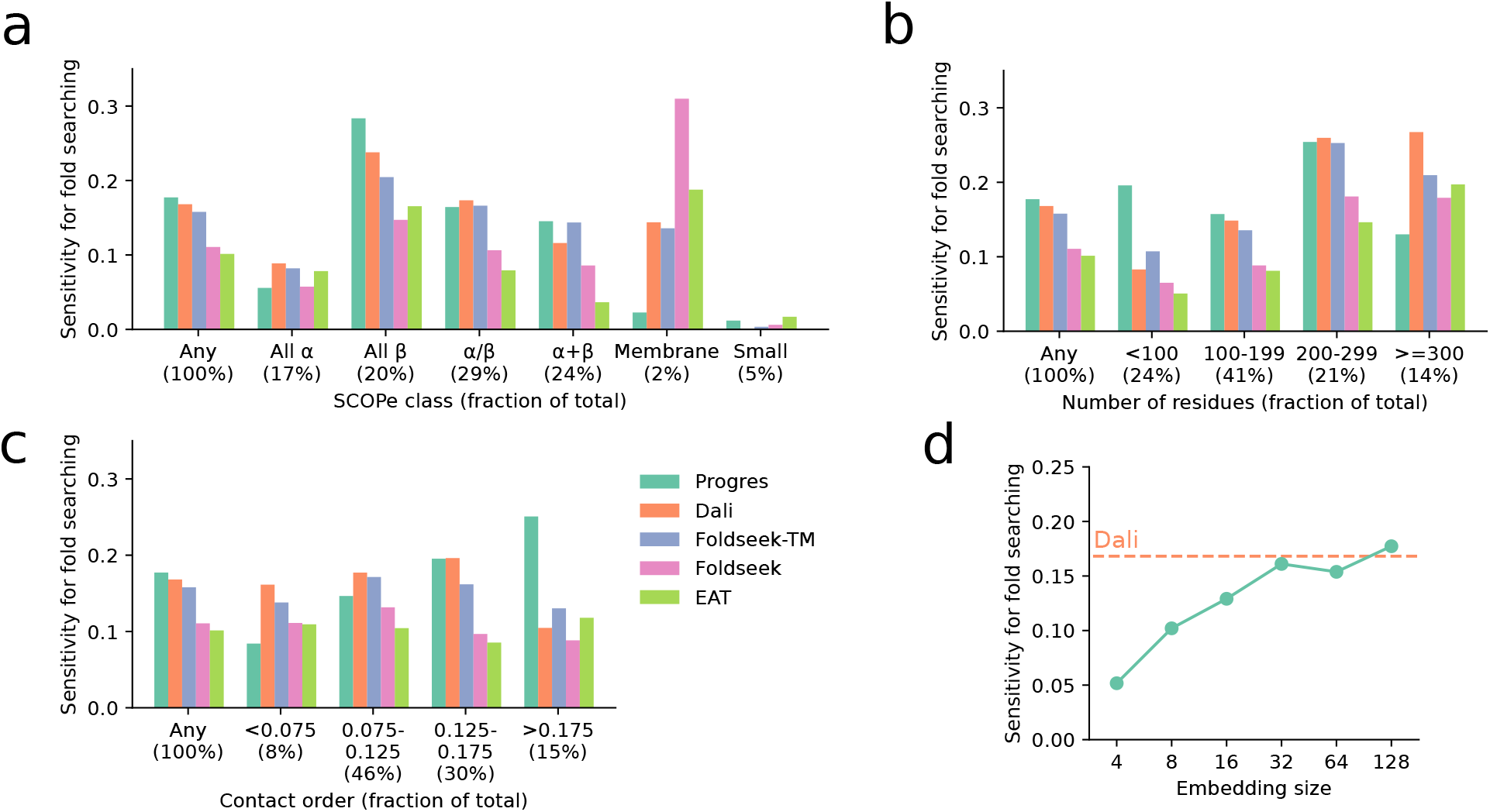
Model performance on different protein types. In each case the “Any” category is the same as in Table 1. (A) Sensitivity for fold searching by SCOPe class. (B) Sensitivity for fold searching by protein sequence length. (C) Sensitivity for fold searching by contact order, a measure of the sequence separation of contacting residues. (D) Sensitivity for fold searching across different embedding sizes. A model was trained from scratch for each embedding size. See Table S1 for further ablations.

For searching a single structure against SCOPe on CPU the model is faster than Foldseek with most run time in Python module loading. For example, going from 1 to 100 query structures increases run time from 1.3 s to 2.4 s. When searching with multiple structures, most run time is in generating the query structure embeddings. Consequently, the speed benefits of the method arise when searching a structure or structures against the pre-computed embeddings of a huge database such as the AlphaFold database [10, 6, 7]. The recent TED study split the whole AlphaFold database into domains using a consensus-based approach [45]. We embed the TED domains clustered at 50% sequence identity and use FAISS [46] to considerably speed up the search time against the resultant database of 53 million structures. This allows a search time of a tenth of a second per query on CPU, after an initial data loading time of around a minute. Since we search exhaustively with FAISS, the results are not changed, though the approximate score calculation means the similarity score does vary slightly from the exact value. For the SCOPe test set used above, the mean difference between FAISS and exact similarity scores for the top hit is 0.006. As shown in Figure S2, the best TM-align score to the query among the top 5 hits has a mean of 0.80 across the SCOPe test set, with 94% being over 0.5. This indicates that searching is accurate even when using a large database.

Figure 3 shows 2D t-SNE embeddings [47] of the 128 dimensions of our model embedding. This shows the lower-dimensional protein fold space [48–50] created by our embedding. SCOPe classes tend to cluster together, with α+β folds appearing between the all-α and all-β folds which show little overlap. There is a clear protein size gradient across the t-SNE embedding. A t-SNE embedding for the AlphaFold database TED domains compared to ECOD [22] and the AlphaFold 21 organisms set [17, 51] shows the volume of new structural information available in the AlphaFold database. The Progres score between two embeddings is the cosine similarity score normalised to run between 0 and 1, with 1 indicating identical embeddings. As shown in Figure 3E a Progres score of 0.8 indicates that two proteins share the same fold, analogous to a TM-align score of 0.5.

**Figure 3.**
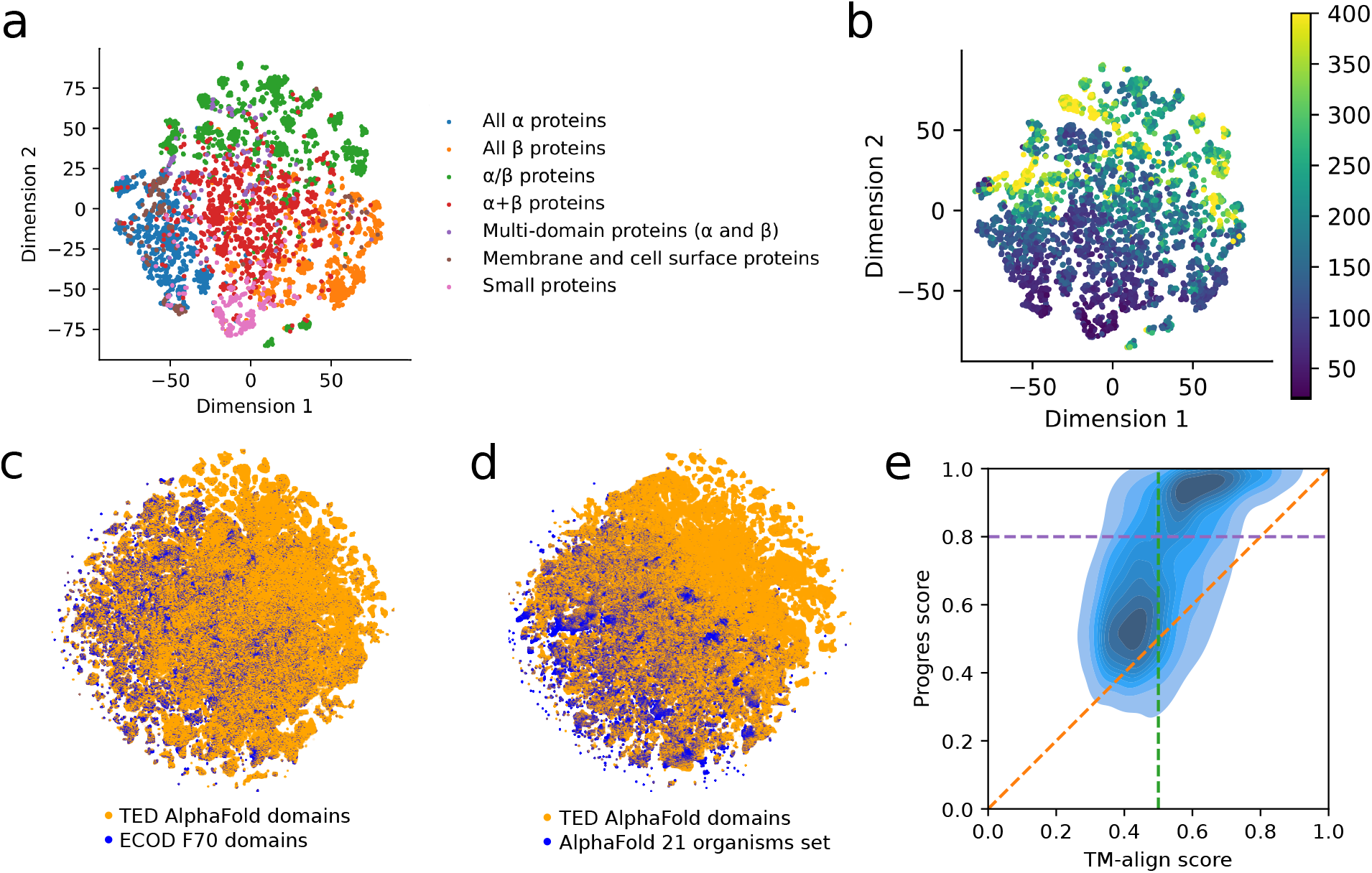
Exploring Progres embeddings. (A) 2D t-SNE embedding of the 128 dimensions of our model embedding for the Astral set of SCOPe domains clustered at 40% sequence identity (15,177 domains). The domains are coloured by SCOPe class. t-SNE was carried out using a perplexity value of 30. (B) The same data coloured by number of residues in the domain. The median length of domains is 149 residues. For colouring, the maximum number of residues in a domain is treated as 400. (C) 2D t-SNE of the AlphaFold database TED domains [45] clustered at 50% sequence identity and the ECOD F70 set of domains in the PDB [22]. 5m (9%) of the TED domains are chosen randomly for the t-SNE for computational reasons. (D) A similar comparison of the TED domains to the AlphaFold 21 model organisms set [17, 51]. (E) Comparison of Progres score to TM-align score. For each of the 400 domains in the test set the top 200 matches in the Astral 40% sequence identity set according to TM-align are considered. The Pearson correlation coefficient is 0.60. The green line shows the TM-align score threshold of 0.5 indicating the same fold. The purple line shows the Progres score threshold of 0.8 indicating the same fold.

## Discussion

The model presented here is trained and validated on protein domains; due to the domain-specific nature of the training it is not expected to work without modification on protein chains containing multiple domains, long disordered regions or complexes. Fortunately, there are a number of tools such as Chainsaw [52], Merizo [53] and SWORD2 [54] that can split query structures into domains. We integrate Chainsaw into Progres to allow automated splitting of query structures into domains, with each domain then searched separately. This can overcome issues that arise from searching with multiple domains at the same time, such as missing related proteins due to differing orientations of the domains. Splitting with Chainsaw takes a few seconds per query. The web server allows visualisation of query and hit domains. As shown in Figure S3 the Progres embeddings are fairly robust to truncating residues from the termini, with truncations of 20 residues giving an embedding with a similarity of 0.8 to the full length domain embedding for 89% of domains with 200-299 residues. This means that minor inaccuracies in predicting domain boundaries are unlikely to cause a problem.

One issue with supervised learning on domains is whether performance drops when searching with domains that the model has not seen anything similar to during training. We trained an identical model on a different dataset where 200 domains were used for testing and domains were removed from the training set if they were from the same SCOPe superfamily as any of the testing domains. The fold, superfamily and family sensitivities analogous to Table 1 are 0.190, 0.383 and 0.546 respectively. This indicates similar performance at finding distantly related folds, the main use of structure searching over sequence searching, though there is a drop in performance at finding closely-related domains.

Aside from searching for similar structures, an accurate protein structure embedding has a number of uses. Fast protein comparison is useful for clustering large sets of structures, for example to identify novel folds in the AlphaFold database [6, 7, 45]. The embedding of a structure is just a set of numbers, and therefore can be targeted by differentiable approaches for applications like protein design. A decoder could be trained to generate structures from the embedding space [55, 56], and a diffusion model to move through the embedding space. Properties of proteins such as evolution [57], topological classification [58], the completeness of protein fold space [59], the continuity of fold space [60], function [61] and dynamics could also be explored in the context of the low-dimensional fold space. Structure embeddings could also be used to identify regions of unknown density in cryo-electron tomography studies. We believe that the extremely fast pairwise comparison allowed by structural embeddings is an effective way to take advantage of the opportunities provided by the million-structure era.

## Methods

### Training

Structures in the Astral 2.08 95% sequence identity set including discontinuous domains were used for training [62]. We chose 400 domains randomly from the Astral 2.08 40% sequence identity set to use as a test set (see below) and another 200 domains to use as a validation set to monitor training. We removed domains with 30% or greater sequence identity to these 600 domains using MMseqs2 [43], and also removed domains with fewer than 20 or more than 500 residues. This left 30,549 domains in 4,862 families for training.

mmCIF files were downloaded and processed with Biopython [63]. Some processing was also carried out with BioStructures.jl [64]. Cα atoms were extracted for the residues corresponding to the domain. Each Cα atom is treated as a node with the following features: number of Cα atoms within 10 Å divided by the largest such number in the protein, whether the Cα atom is at the N-terminus, whether the Cα atom is at the C-terminus, and a 64-dimensional sinusoidal positional encoding for the residue number in the domain [42].

PyTorch was used for training [65]. The neural network architecture was similar to the E(n)-equivariant GNN in Satorras et al. 2021 [20]. We used a PyTorch implementation (https://github.com/lucidrains/egnn-pytorch) and a configuration similar to the molecular data prediction task, i.e. not updating the particle position. In this case the model is analogous to a standard GNN with relative squared norms inputted to the edge operation [20]. Edges are sparse and are between Cα atoms within 10 Å of each other. 6 such layers with residual connections are preceded by a one-layer multilayer perceptron (MLP) acting on node features and followed by a two-layer MLP acting on node features. Node features are then sum-pooled and a two-layer MLP generates the output embedding, which is normalised. Each hidden layer has 128 dimensions and uses the Swish/SiLU activation function [66], apart from the edge MLP in the GNN which has a hidden layer with 256 dimensions and 64-dimensional output. The final embedding has 128 dimensions.

Supervised contrastive learning [23] is used for training. Each epoch cycles over the 4,862 training families. For each family, 5 other families are chosen randomly. For each of these 6 families, 6 domains from the family present in the training set are chosen randomly. If there are fewer than 6 domains in the family, duplicates are added to give 6. This set of 36 domains with 6 unique labels is embedded with the model and the embeddings are used to calculate the supervised contrastive loss with a temperature of 0.1 [23]. During training only, Gaussian noise with variance 1.0 Å is added to the x, y and z coordinates of each Cα atom. Training was carried out with the Adam optimiser [67] with learning rate 5 × 10^−5^ and weight decay 1 × 10^−16^. Each set of 36 domains was treated as one batch. Training was stopped after 500 epochs and the epoch with the best family sensitivity on the validation set was used as the final model. Training took around a week on one RTX A6000 GPU.

### Testing

For testing a similar approach to Foldseek was adopted [19]. The 15,177 Astral 2.08 40% sequence identity set domains were embedded with the model. The embeddings are stored as Float16 to reduce the size of large databases on disk, but this has no effect on search performance as shown in Table S1. 400 of these domains were chosen randomly and held out of the training data as described previously. Like Foldseek, we only chose domains with at least one other family, superfamily and fold member. For each of these 400 domains, the cosine similarity of embeddings to each of the 15,177 domains was calculated and the domains ranked by similarity with the query domain included. For each domain, we measured the fraction of TPs detected up to the first incorrect fold detected. TPs are same family in the case of family-level recognition, same superfamily and not same family in the case of superfamily-level recognition, and same fold and not same superfamily in the case of fold-level recognition. We also report the mean TM-align score and the fraction of hits with the same fold for the top 20 hits for each query.

All CPU methods were run on an Intel i9-10980XE CPU and with 256 GB RAM. Progres, Foldseek and MMseqs2 were run on 16 threads. The GPU methods were run on a RTX A6000 GPU. Progres was run with PyTorch 1.11. Foldseek version 8.ef4e960 was used. For TM-align we used the fast mode, which has similar performance to the normal mode [19]. For 3D-SURFER the neural network model and mainchain atoms were used. EAT was run with the “–use tucker 1” flag. ESM-2 embeddings used the esm2 t36 3B UR50D model which has a 2560-dimensional embedding. The mean of the per-residue representations was normalised and comparison between sequences was carried out with cosine similarity. For MMseqs2, easy-search with a sensitivity of 7.5 was used.

For contact order, all residue pairs with Cβ atoms (Cα for glycine) within 8 Å are considered. The contact order of a structure is then defined as

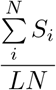

where *S*_*i*_ is the sequence separation of the residues in contacting pair *i, N* is the number of contacting pairs and *L* is the sequence length of the protein.

## Databases

The AlphaFold database domain embeddings were prepared from the TED set of domains [45] using cluster representatives from clustering at 50% sequence identity. Clustering was carried out with MMseqs2 using the command “mmseqs easy-cluster ted 100.fasta clusterRes tmp –min-seq-id 0.5 –c 0.9 –cov-mode 5 -s 7.5”. This gave 53,344,209 clusters, fewer than the TED analysis due to the use of easy-cluster over easy-linclust. The FAISS [46] index was prepared using “IndexFlatIP(128)”, which carries out exhaustive searching using the same cosine similarity as Progres. Query structures may be automatically split into domains before searching using Chainsaw [52], with each domain searched separately.

## Availability

A Python package allowing structure searching and generation of pre-embedded databases is available along with datasets, training scripts and benchmarking scripts under a permissive license at https://github.com/greener-group/progres. The trained model and pre-embedded databases are available at https://zenodo.org/record/7782088. A web server for searching is available at https://progres.mrc-lmb.cam.ac.uk.

## Acknowledgements

We thank the UCL Bioinformatics Group for useful discussions and Jake Grimmett, Toby Darling and Ivan Clayson for help with high-performance computing and web server deployment. KJ is a member of the group of Sjors Scheres. This work was supported by the Medical Research Council, as part of United Kingdom Research and Innovation (also known as UK Research and Innovation) [MC UP 1201/33 to JGG and MC UP A025-1013 to SS]. For the purpose of open access, the MRC Laboratory of Molecular Biology has applied a CC BY public copyright licence to any Author Accepted Manuscript version arising.

## Supplementary Data

**Table S1.** Model ablations. In each case a model is trained from scratch and the same testing procedure is used as in Table 1. No ablation is the model presented in the results with 128 embedding dimensions and 6 layers. The embedding dimension ablation is shown visually in Figure 2D. Removing the τ angle node feature means the network is E(3)-invariant not SE(3)-invariant. Superfamily holdout validation is described in the discussion.

| Ablation | Fold | Superfamily | Family |
| --- | --- | --- | --- |
| No ablation | 0.177 | 0.706 | 0.877 |
| 64 embedding dimensions | 0.154 | 0.673 | 0.864 |
| 32 embedding dimensions | 0.161 | 0.699 | 0.884 |
| 16 embedding dimensions | 0.129 | 0.654 | 0.850 |
| 8 embedding dimensions | 0.102 | 0.491 | 0.696 |
| 4 embedding dimensions | 0.052 | 0.181 | 0.353 |
| 12 layers | 0.150 | 0.709 | 0.902 |
| 4 layers | 0.148 | 0.645 | 0.841 |
| No $\tau$ angle node feature | 0.159 | 0.690 | 0.869 |
| No Gaussian noise | 0.072 | 0.516 | 0.738 |
| No sinusoidal positional encoding | 0.052 | 0.402 | 0.642 |
| Float32 database for searching | 0.177 | 0.706 | 0.877 |
| Superfamily holdout validation | 0.190 | 0.383 | 0.546 |

**Figure S1.**
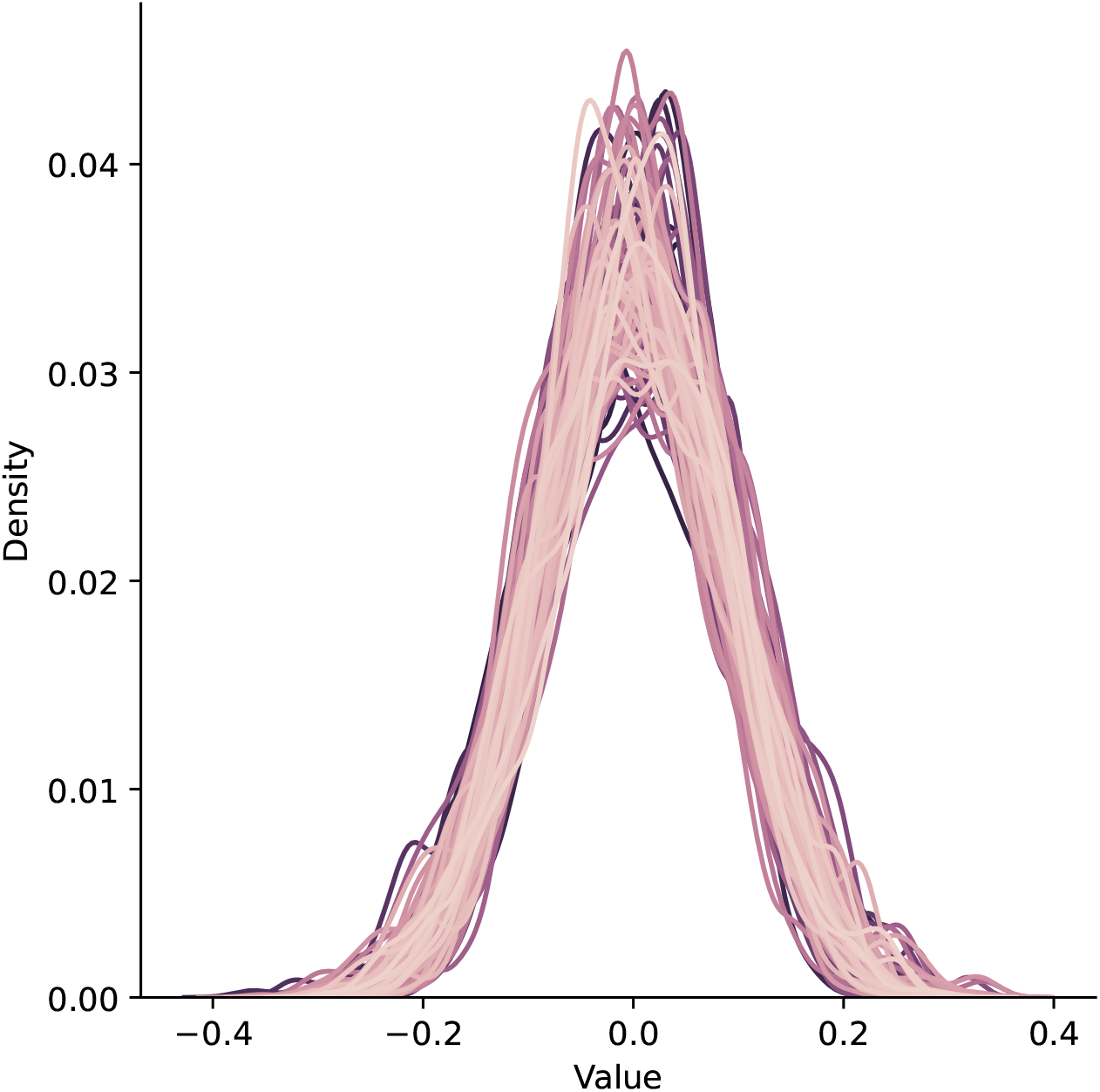
Values of embedding dimensions. Each protein in the Astral 40% sequence identity set is embedded with Progres and the distribution of values in each of the 128 dimensions is shown.

**Figure S2.**
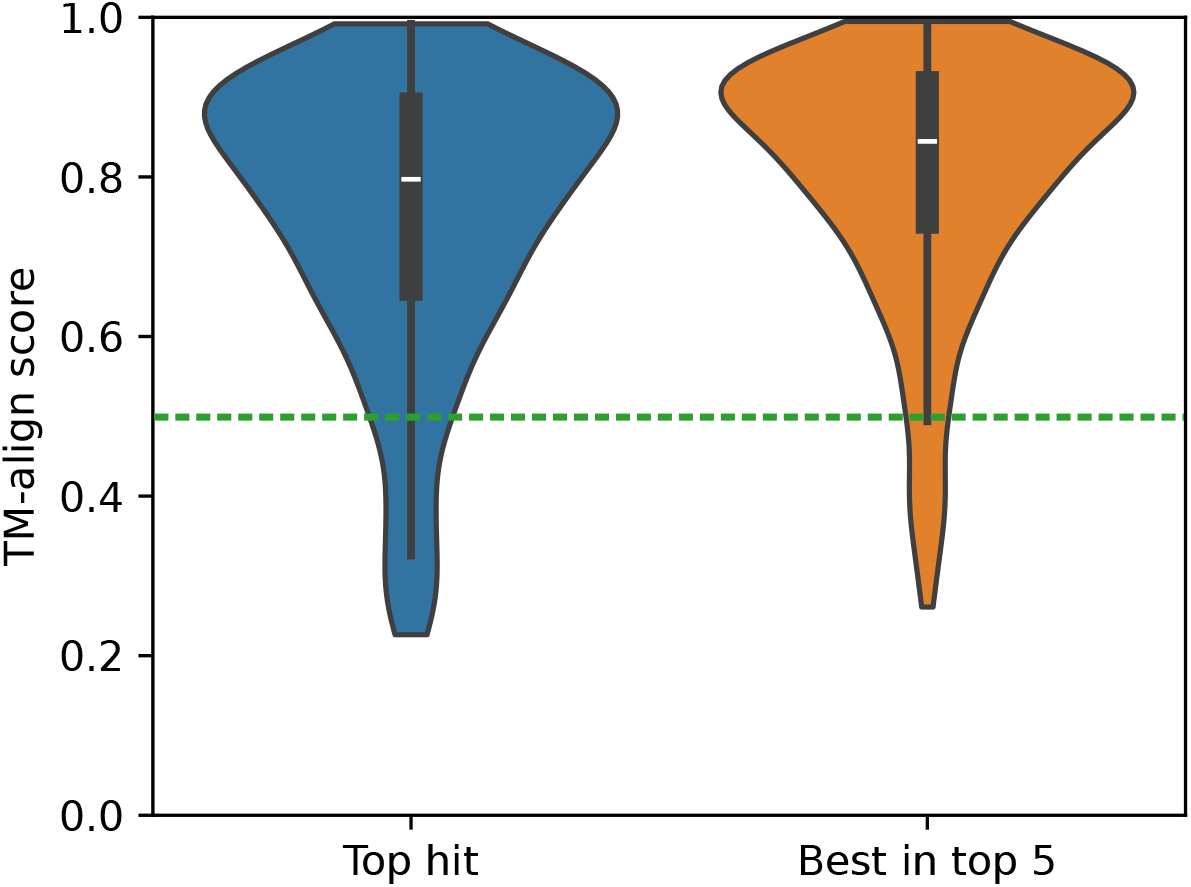
Searching SCOPe domains against the AlphaFold database. Each of the 400 domains in the test set is searched against the AlphaFold database TED domains clustered at 50% sequence identity (53 million domains) and the TM-align score between the top hit and the query is calculated. This gives a mean TM-align score of 0.75 with 90% above 0.5. The highest TM-align score among the top 5 hits is also calculated and gives a mean of 0.80 with 94% above 0.5.

**Figure S3.**
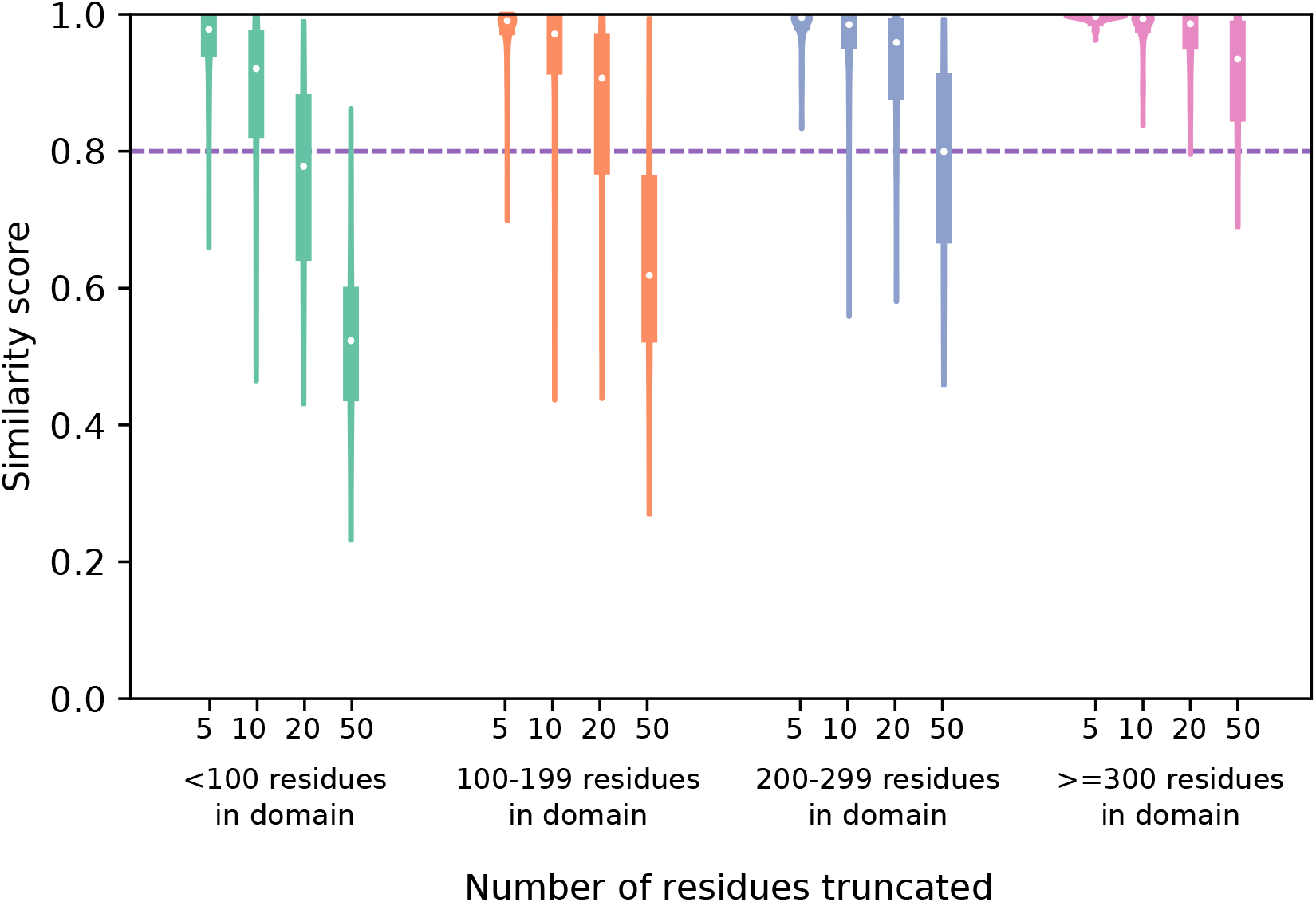
The effect of truncating domains on Progres embeddings. For each of the 400 domains in the SCOPe test set a number of residues were removed from the N-terminus or the C-terminus and the truncated domain was embedded. The Progres similarity score to the full length domain was then computed. The results are categorised by the number of residues in the full length domain. The line shows a similarity score of 0.8, indicating the same fold (see Figure 3E).

